# golgi: open-source software for automated nerve model generation and recruitment simulation

**DOI:** 10.64898/2026.07.10.737846

**Authors:** David Lung, Yuting Jia, Andrea Moro, Matteo Fachino, Max Haberbusch

## Abstract

*golgi* is an open-source platform that takes a peripheral nerve from image to stimulated fiber population through a single graphical interface, with an equivalent scriptable Python API and command-line interface for batch and high-performance use. It integrates promptable image segmentation, automated multi-region tetrahedral meshing, anisotropic finite-element solution of the extracellular field with an explicit perineurium contact impedance, generation of realistic fiber populations and their three-dimensional trajectories, and biophysical activation thresholds through interchangeable backends— NEURON (via PyFibers) and a GPU-accelerated surrogate (AxonML). Every study exports as an integrity-hashed bundle whose image-to-recruitment provenance is verifiable byte-for-byte. *golgi* lowers the barrier to in-silico peripheral nerve stimulation modeling for experimentalists and clinicians, using a fully open finite-element stack with no commercial dependencies.

## 1. Motivation and significance

Electrical stimulation of peripheral nerves treats a growing range of conditions, from epilepsy to inflammatory and cardiovascular disease. Deciding where to place an electrode and how to shape the stimulus increasingly relies on computational models that couple a volume-conductor description of the tissue—most often solved by the finite-element method (FEM)—to biophysical cable models of individual axons [9, 8]. The extracellular potential computed along each fiber drives a multi-compartment membrane model, and the second spatial derivative of that potential (the activating function) predicts where the fiber is depolarized. Combined with realistic fascicular geometry, this lets recruitment and selectivity be estimated *in silico*.

The established open tools for this workflow are powerful but *code-first* : they assume substantial modeling expertise, and several depend on a commercial finite-element solver. ASCENT [2], the de-facto standard pipeline for vagus-nerve stimulation, drives its field solution through COMSOL Multiphysics; NRV [3] is a scriptable Python framework. Both require the user to operate the model through code and configuration files. This places anatomically realistic nerve modeling out of reach for many of the experimentalists and clinicians who could most benefit from it, and the dependence on proprietary software is a barrier to open, reproducible science.

*golgi* was built to remove that barrier. It exposes the entire image-to-recruitment pipeline through a graphical user interface (GUI) usable without programming, while mirroring every operation in a scriptable Python API and a command-line interface (CLI) for batch and high-performance-computing use. It builds anatomically realistic, multifascicular nerve models— directly from segmented images, imported surfaces, or label masks—meshes and solves them with a fully open FEM stack (FEniCSx/DOLFINx [4]), generates realistic fiber populations with straight or physically curved three-dimensional trajectories, and computes biophysical thresholds through interchangeable simulation backends (NEURON [5] via PyFibers [7], and the AxonML GPU surrogate [6]). Uniquely, *golgi* exports each study as an integrity-hashed, self-contained bundle whose finite-element-to-recruitment provenance is verifiable byte-for-byte with a single command. A companion paper [1] describes the underlying methods, validates *golgi* against three independent recruitment benchmarks, and uses it to study branch-selective vagus-nerve stimulation in reconstructed three-dimensional rabbit and human nerves; the present paper describes the software itself.

**Table 1.**
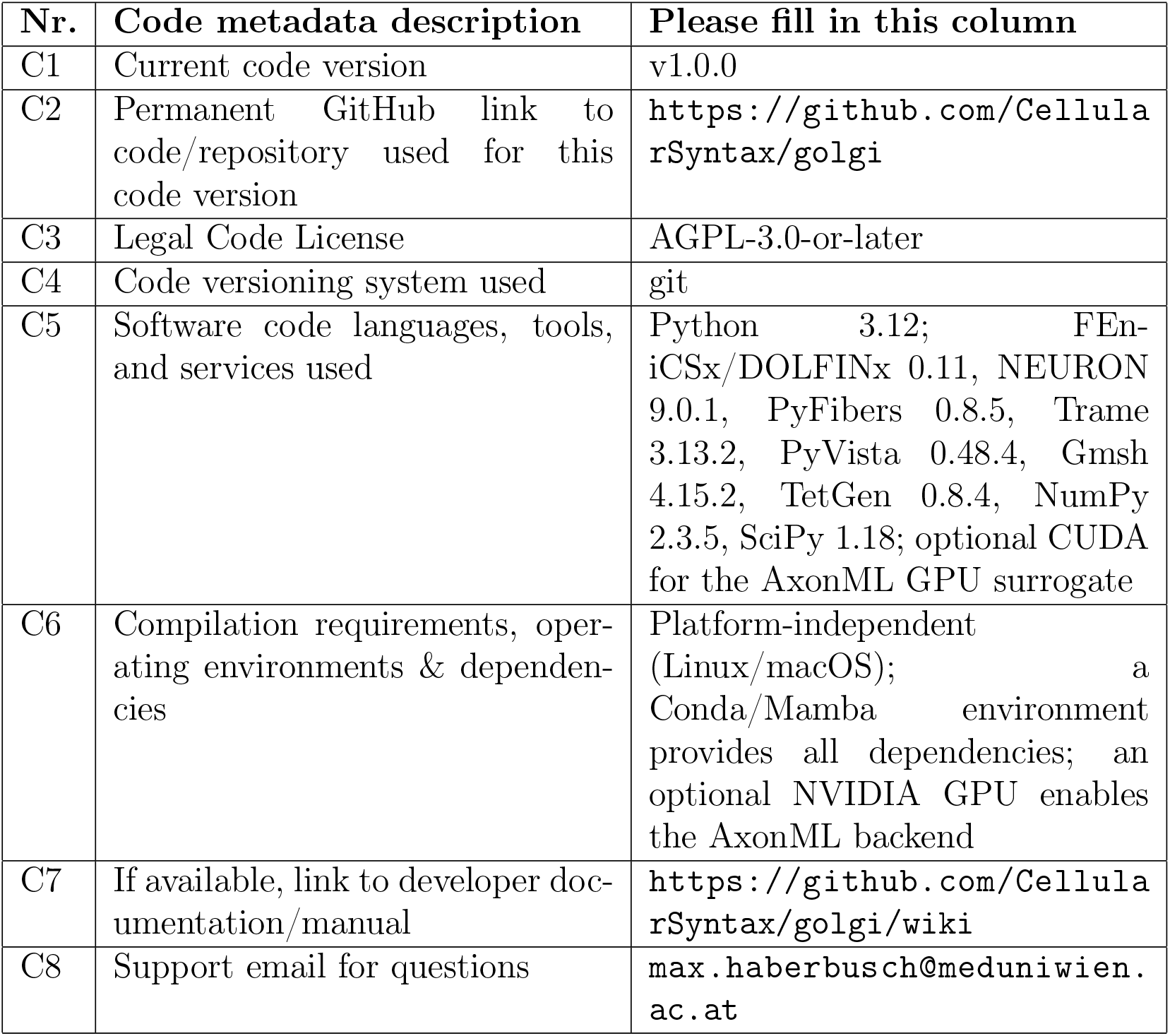
Code metadata.

## 2. Software description

### 2.1. Software architecture

*golgi* is organized as a unified computational pipeline (Fig. 1) that begins with image or geometric data and proceeds through multi-region mesh generation, finite-element field construction, fiber-population and trajectory generation, biophysical fiber simulation, and downstream analysis. A single persistent object, `golgi.Study`, holds the complete state of a study (geometry, mesh, electrode designs, fields, fiber populations, simulation results) and is the common substrate for all three interfaces:

- an interactive, browser-based **GUI** built with Trame for point-and-click operation (Fig. 3-4);
- a headless **Python API** (`golgi.Study`) that mirrors every GUI action for scripting and batch studies;
- a **command-line interface** for cluster and continuous-integration use.

**Figure 1.**
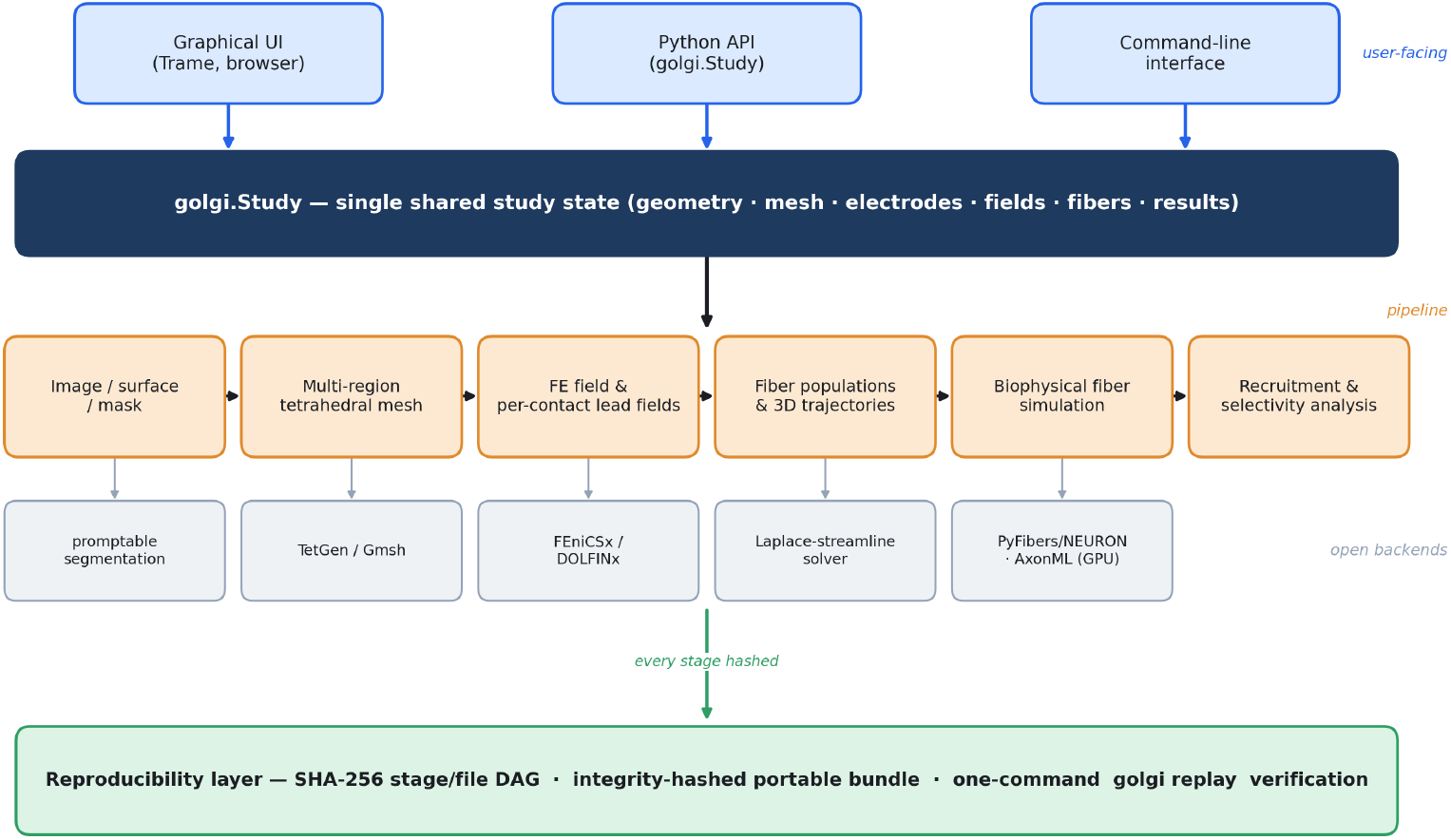
High-level architecture of *golgi*. The three user-facing interfaces (graphical UI, Python API, and CLI) all act on a single shared `golgi.Study` state. The image-to-recruitment pipeline runs on open backends—promptable segmentation, TetGen/Gmsh meshing, the FEniCSx/DOLFINx field solver, a Laplace-streamline trajectory solver, and PyFibers/NEURON or the AxonML GPU surrogate for fiber simulation—and a reproducibility layer hashes every stage into an integrity-verified, portable bundle.

Every GUI action maps onto an API call on the same `Study`, so a non-programming user can complete a full study graphically while an advanced user can script identical studies or offload them to a scheduler, with results interchangeable across modes. Computationally heavy stages run as managed jobs (local subprocesses or cluster submissions), keeping the interface responsive. The field solution uses FEniCSx/DOLFINx; fiber simulation dispatches to a validated NEURON backend (PyFibers) or a GPU surrogate (AxonML) behind a common interface. A reproducibility layer wraps the whole pipeline: each stage records a directed acyclic graph (DAG) of inputs and outputs with SHA-256 hashes, enabling a one-command, byte-for-byte replay (Section 2.2).

### 2.2. Software functionalities

*golgi* ‘s major functionalities, each available identically through the GUI and API, are:

1. **Geometry creation**. Nerve geometry can be created three ways: by segmenting a volumetric image (e.g. a micro-CT stack) with an integrated promptable segmentation engine guided by point and box prompts, followed by interactive mask refinement (per-class reassignment, brush add/erase) [12, 13]; by importing a surface mesh (STL/NAS/OBJ); or by importing label masks (e.g. from histology). Segmented or imported cross-sections are reconstructed into a three-dimensional nerve by extrusion of a representative section or, for genuine three-dimensional samples, from the full stack.
2. **Multi-region meshing**. An automated mesher (TetGen/Gmsh [11]) builds a conforming tetrahedral mesh comprising endoneurium, perineurium, epineurium, the electrode and its insulating cuff, and the surrounding bath, with interactive mesh-quality visualization.
3. **Electrode design**. An interactive cuff designer places and parameterizes electrodes—from bipolar and multi-contact ring cuffs to longitudinal intrafascicular electrodes (LIFE)—and automatically fits the cuff to the local longitudinal axis of the nerve, with rotational and longitudinal position sweeps.
4. **Finite-element field**. *golgi* solves the quasi-static volume-conductor problem with FEniCSx/DOLFINx, using anisotropic endoneurial conductivity and an explicit perineurium contact impedance [10]. By superposition, a per-contact lead field is computed once and reused, so arbitrary multi-contact configurations and current-steering combinations are evaluated without re-solving the field.
5. **Fiber populations and trajectories**. Each fascicle is populated with fibers drawn from species- and nerve-specific diameter and fibertype statistics, mixing myelinated (A- and B-type) and unmyelinated (C-type) fibers. Trajectories are generated either as straight axial extrusions or, for three-dimensional and branching geometries, as smooth curved paths obtained by integrating streamlines of a quasi-static field within the endoneurial domain.
6. **Biophysical simulation and recruitment**. Activation thresholds and responses are computed with validated axon models (e.g. MRG, Sundt, Tigerholm) through interchangeable backends—NEURON (PyFibers) as the gold standard and the AxonML GPU surrogate [6] for high-throughput sweeps. Thresholds are found by bisection on configurable mono- or biphasic waveforms; recruitment and fascicular/branch selectivity follow from amplitude sweeps and current steering.
7. **Reproducible export**. Any study exports as a single self-contained bundle (Fig. 2) whose manifest records the pipeline DAG with a SHA-256 hash of every input, output, and stage, together with the exact software version and a frozen dependency list. A single `golgi replay` command re-hashes a received bundle and verifies byte-for-byte that it is intact and matches its manifest, or flags the first stage that differs.
8. **Visualization**. Publication-quality three-dimensional renderings of the nerve, electrode, fields, and recruited fibers are produced directly from the pipeline.

**Figure 2.**
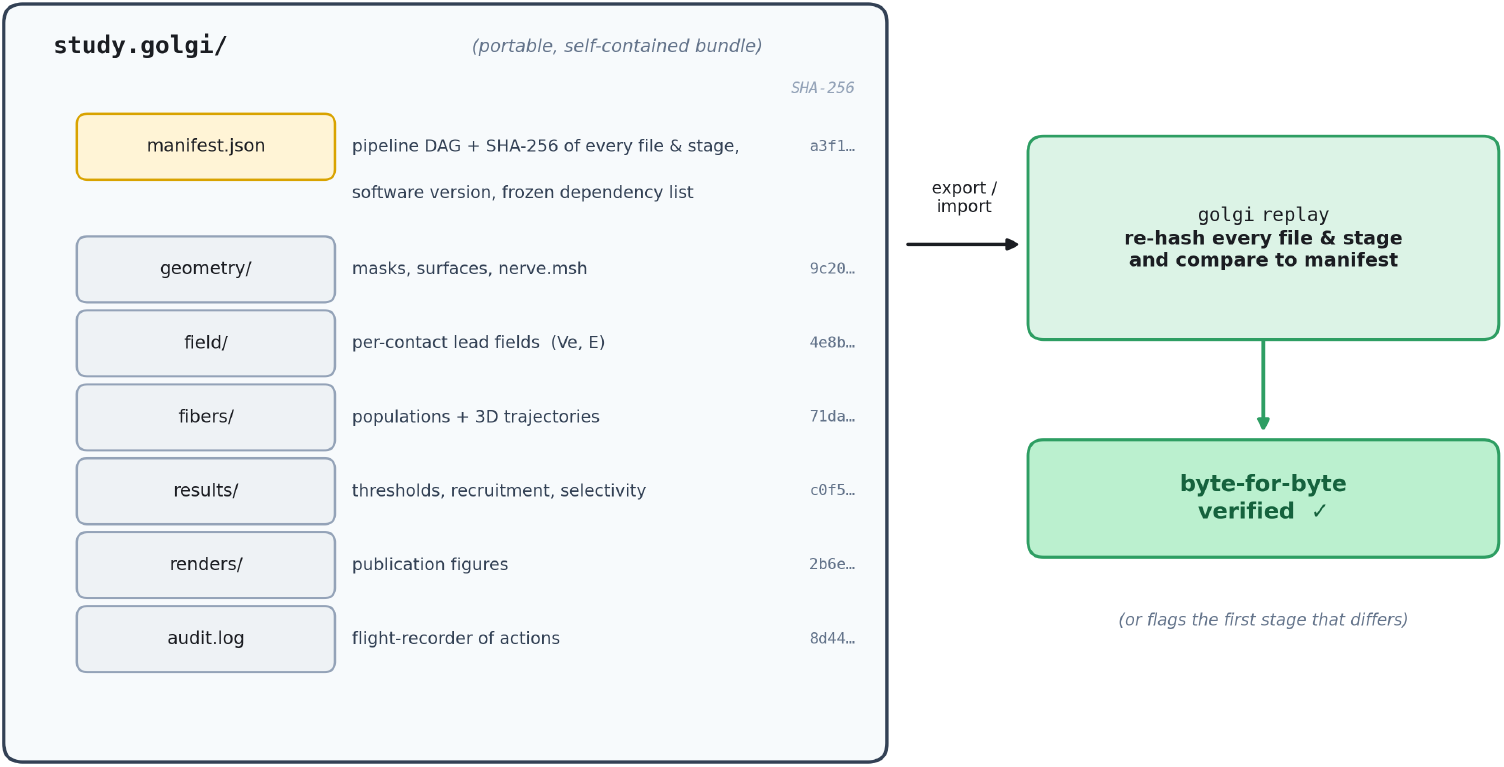
Reproducible study bundle. Every study exports as a portable .golgi bundle: a manifest.json records the pipeline directed acyclic graph and a SHA-256 hash of every file and stage (plus the software version and a frozen dependency list), alongside the geometry, fields, fibers, results, renders, and a flight-recorder audit log. On import, golgi replay re-hashes the bundle and verifies it byte-for-byte against the manifest—or flags the first stage that differs.

**Figure 3.**
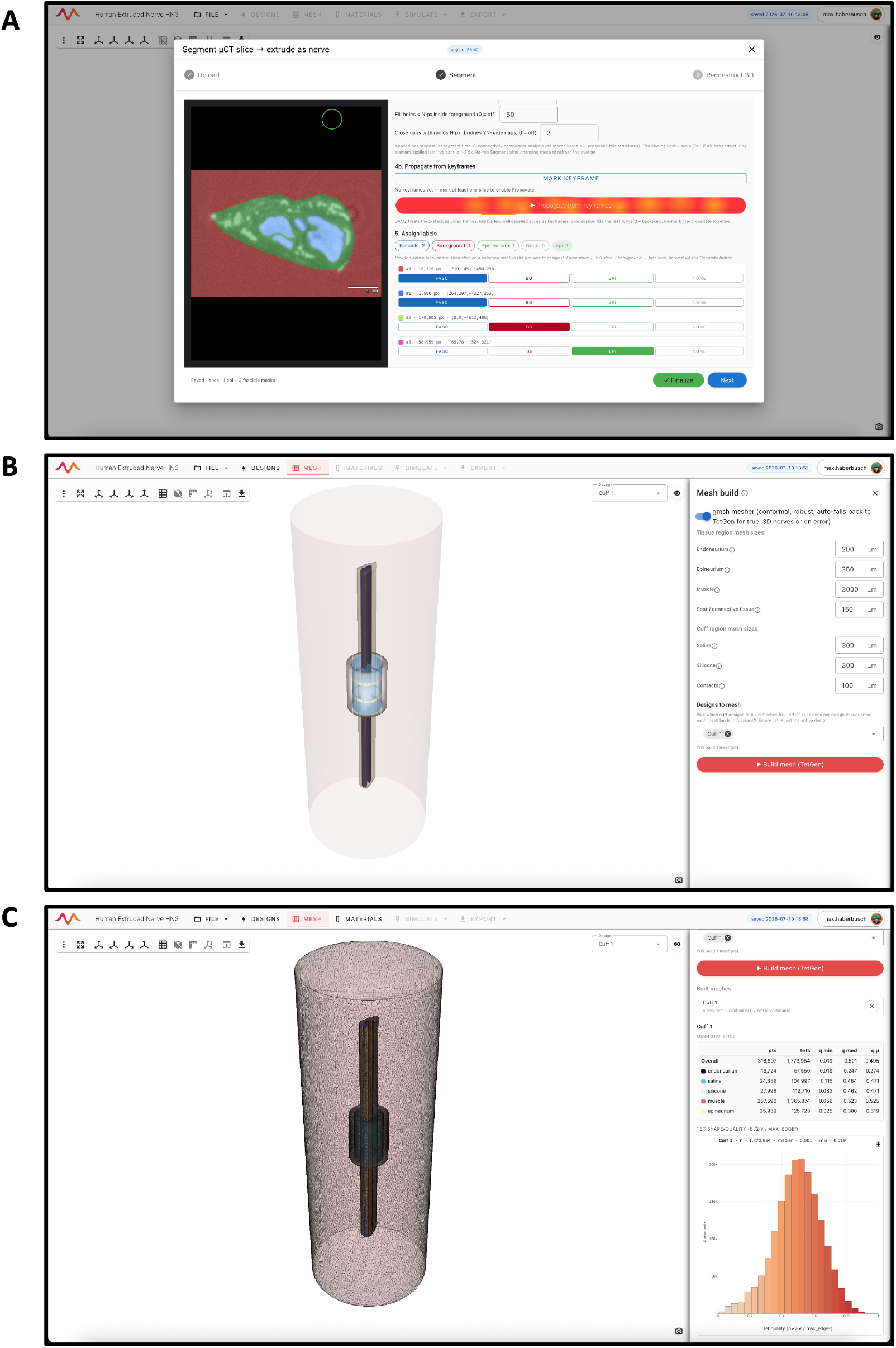
The golgi graphical user interface at three representative pipeline stages: (A) segmented micro-CT image, (B) extruded nerve and fitted cuff electrode, (C) multi-region mesh from the geometry defined in the previous steps.

### 2.3. Sample code

The headless API mirrors the GUI; a complete study is a short script:

~~~
import golgi
s = golgi.Study.create(“vagus_study”)
s.import_geometry(“vagus_microCT.tif”) # image/surface/mask
s.segment(prompt=“fascicle”)# promptable segmentation
s.set_electrodes([dict(kind=“ring-array”, n_rows=4, n_cols=5)]) # interactive cuff
s.set_mesh(perineurium_ci=True) # regions + contact impedance
s.run_mesh()
s.set_fiber_population(preset=“cervical_vagus_human”)
s.set_fiber_seed(n_fibers=600, fiber_method=“streamlines”)
s.run_fem() # anisotropic FEM + lead fields
s.run_sweep(backend=“pyfibers”) # thresholds / recruitment
s.export_bundle(“vagus_study.golgi”) # hashed, replayable bundle
~~~

## 3. Illustrative examples

Figures 3–4 show the GUI at six representative pipeline stages. **(1) Image segmentation:** a micro-CT slice of a cervical vagus nerve is segmented into fascicles, epineurium, and background using the promptable SAM2 engine, with the endoneurial compartment taken as the fascicle interior (Fig. 3A). **(2) Geometry generation & electrode fitting:** the segmented cross-section is extruded into a three-dimensional nerve, and a multi-contact cuff is placed and auto-fitted around the nerve at its axial center (Fig. 3B). **(3) Mesh generation:** a multi-region tetrahedral mesh is built with TetGen, with per-region element counts and tetrahedron shape-quality reported for inspection (Fig. 3C). **(4) Conductivity assignment:** quasi-static conductivities are assigned to each tissue and electrode region—optionally via a Cole–Cole evaluator at the stimulation frequency—and passed to the FEM solver (Fig. 4A). **(5) Simulation:** the solved extracellular potential *V*_*e*_ is inspected through four linked views: a three-dimensional rendering of *V*_*e*_ on the nerve with the in-plane current streamlines around the active contact and the fiber trajectories threaded through the endoneurium; a transverse *V*_*e*_ slice heatmap at a user-selectable axial position *z*; *V*_*e*_ and the axial field *E*_*z*_ = − *∂V*_*e*_*/∂s* sampled along the fibers; and the activation function *f*_*n*_ = *∂*^2^*V*_*e*_*/∂s*^2^, each shown as a per-branch mean *±*1*σ* with an individually selectable fiber (Fig. 4B). **(6) Analysis:** a stimulus waveform is defined and a population simulation is run, yielding a cross-sectional activation map of recruited versus quiescent fibers and the propagating membrane potential *V*_*m*_ along the fibers—all without writing code (Fig. 4C).

**Figure 4.**
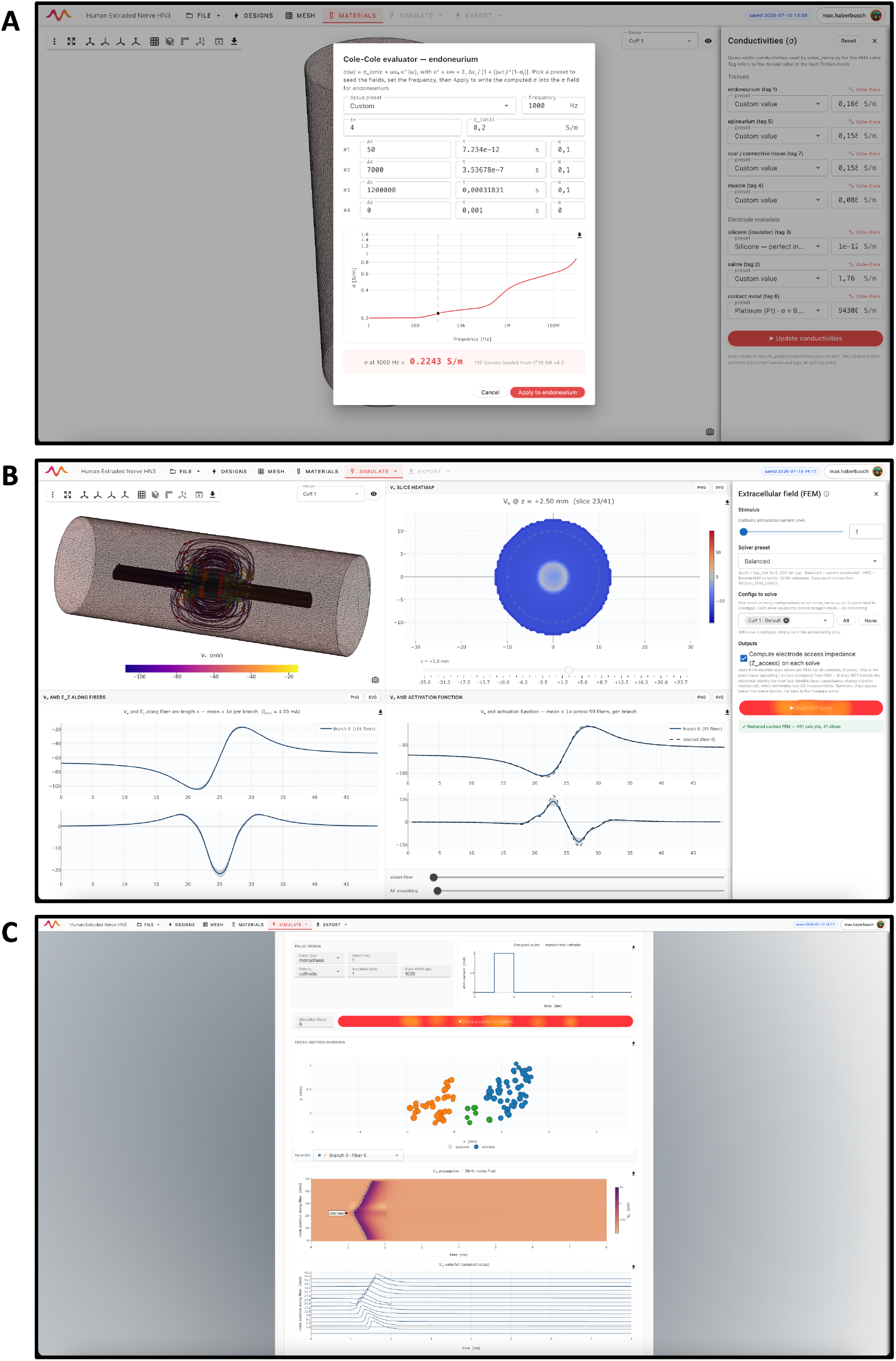
The golgi graphical user interface at three representative pipeline stages: (A) conductivity and material property assignment, (B) FEM-solve overview, (C) nerve activity simulation overview.

The same study is reproducible from the API script in Section 2.3: running it regenerates the mesh, field, fiber population, and recruitment results, and `export_bundle` writes a bundle that any recipient can verify and reuse with `golgi replay vagus_study.golgi`. In the companion study [1], this identical pipeline was applied across a cohort of image-derived swine and human cervical vagus nerves and to reconstructed three-dimensional branching rabbit and human nerves, reproducing physiological fiber-diameter recruitment order and quantifying fascicular and branch selectivity with a multi-contact cuff.

A detailed description and tutorials are available at https://github.com/CellularSyntax/golgi/wiki/Tutorial

## 4. Impact

*golgi* changes who can build anatomically realistic peripheral-nerve-stimulation models and how trustworthy the results are.

First, it **removes the programming and proprietary-software barriers** that have confined realistic nerve modeling to specialist groups. By exposing the complete image-to-recruitment pipeline through a graphical interface backed by a fully open FEM stack, it lets experimentalists and clinicians—not only computational modelers—design electrodes and waveforms, compare cuff configurations, and predict fiber-type-selective recruitment, without writing code or licensing COMSOL. The mirrored API and CLI mean the same study scales seamlessly from an interactive session to a high-performance-computing batch, so exploratory and production use share one tool and one set of results.

Second, it **enables research questions that straight-fiber, cross-section-only tools cannot address**. Because *golgi* reconstructs genuine three-dimensional nerves and generates curved, fascicle-following fiber trajectories through bifurcations, it can quantify *branch-selective* stimulation—steering current from a proximal cuff to a named distal branch and resolving which fiber classes are captured. Using this capability, the companion study [1] finds that whether a clinically important vagal cardiac branch can be selectively engaged depends on its anatomy, a question that previously could not be posed *in silico* on subject-specific geometry.

Third, it **makes nerve-stimulation studies verifiably reproducible**. The integrity-hashed bundle and one-command `golgi replay` provide a byte-for-byte provenance guarantee from finite-element field to recruitment prediction—a property absent from existing tools—which supports sharing, review, and reuse of complete studies.

By combining clinical-grade accessibility, anatomical realism, and verifiable reproducibility in one open package, *golgi* broadens participation in in-silico peripheral nerve stimulation modeling and provides a common, open substrate for method development and cross-study comparison.

## 5. Conclusions

*golgi* is an open-source platform that takes a peripheral nerve from image to stimulated fiber population through a single graphical interface, with an equivalent scriptable API and CLI. It integrates promptable segmentation, automated multi-region meshing, an open anisotropic finite-element field solver with explicit perineurium contact impedance, realistic fiber populations with curved three-dimensional trajectories, and interchangeable fiber backends—NEURON (via PyFibers) and a GPU-accelerated surrogate (AxonML)— and it exports every study as an integrity-hashed, replayable bundle. Validated against established recruitment benchmarks and demonstrated on reconstructed three-dimensional nerves in the companion paper [1], *golgi* lowers the barrier to anatomically realistic, reproducible peripheral nerve stimulation modeling for a broad community of users.

## Data availability and licensing

*golgi* is released as open-source software under the AGPL-3.0-or-later license (code metadata C3/S3); its copyleft terms follow from linking the Gmsh (GPLv2-or-later) and TetGen (AGPL-3.0) meshers and serving a browser GUI. The source code is available on GitHub (https://github.com/CellularSyntax/golgi), and the tagged v1.0.0 release is archived on Zenodo [14]. The data accompanying *golgi* are openly available under the Creative Commons Attribution 4.0 International (CC-BY-4.0) license, distinct from the software license. The micro-CT imaging stacks—including tissue segmentations and three-dimensional reconstructions—of the rabbit [16] and human [17] cervical vagus nerves used to build the three-dimensional demonstration geometries are deposited on Zenodo, and the integrity-hashed *golgi* study bundles behind the reported studies are likewise archived on Zenodo [15]. Each bundle is a self-contained project—geometry, mesh, finite-element lead fields, fiber population, and recruitment—that can be imported into *golgi* and re-verified byte-for-byte with `golgi replay`.

## CRediT authorship contribution statement

**David Lung:** Software, Investigation, Validation, Writing – review & editing. **Yuting Jia:** Software, Investigation, Validation, Writing – review & editing. **Andrea Moro:** Software, Writing – review & editing. **Matteo Fachino:** Software, Writing – review & editing. **Max Haberbusch:** Conceptualization, Methodology, Software, Supervision, Funding acquisition, Writing – original draft, Writing – review & editing.

## Acknowledgements

*golgi* was funded by project VAGUSPEC (Medical University of Vienna, Focus Grant M) and project PREVENT (City of Vienna, Agreement ID H-463816/2023). Yuting Jia gratefully acknowledges support from the China Scholarship Council (CSC, File No. 202408460012). Simulations were performed using the Medical University of Vienna high-performance computing cluster.

## Current executable software version

**Table 2.**
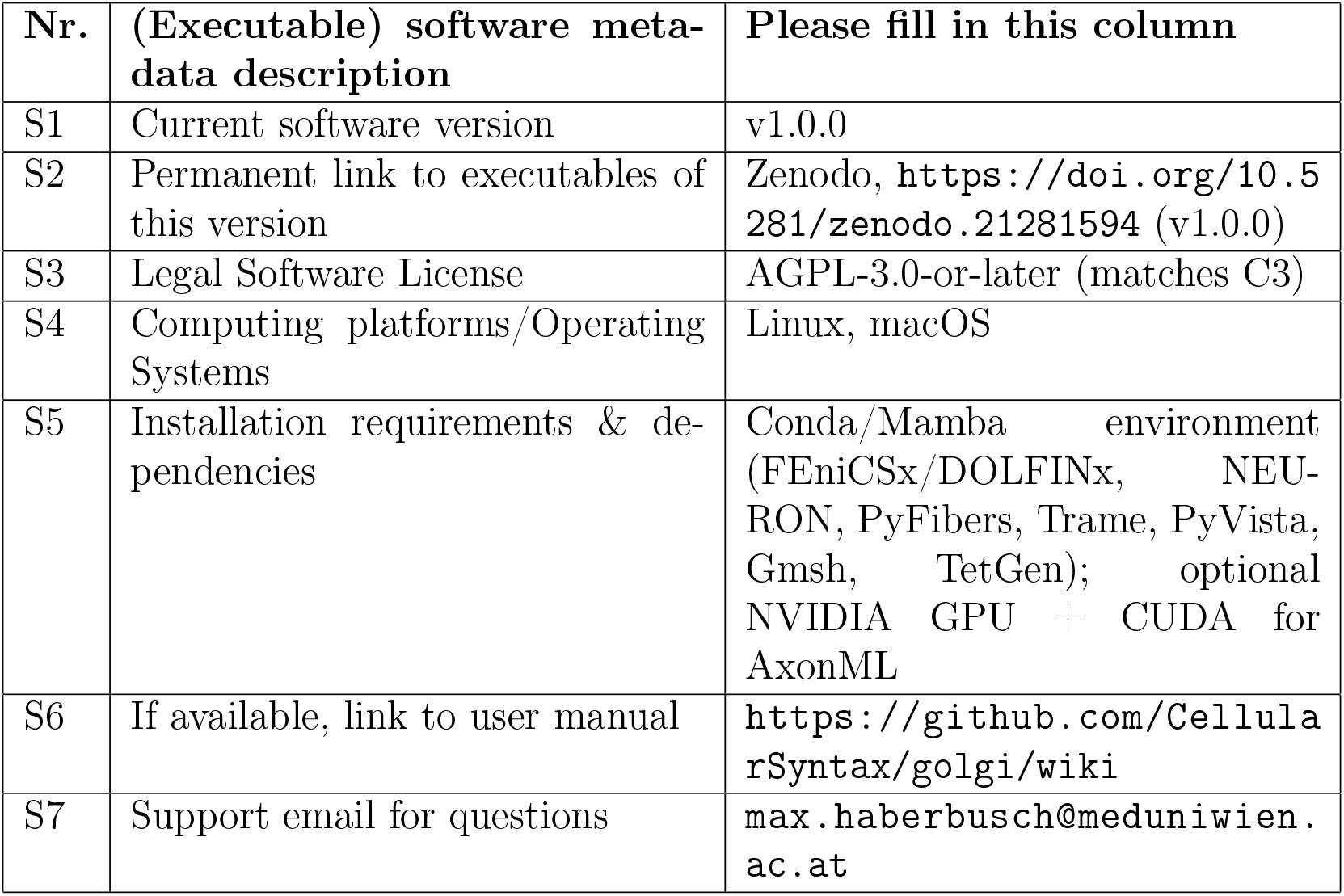
Software metadata.

## Notes

https://github.com/CellularSyntax/golgi/

https://github.com/CellularSyntax/golgi/wiki/

https://doi.org/10.5281/zenodo.21006413

https://doi.org/10.5281/zenodo.21007683

https://doi.org/10.5281/zenodo.21300945

https://doi.org/10.5281/zenodo.21301866

### Competing Interest Statement

The authors have declared no competing interest.

## References

[1] Authors, golgi: an open, reproducible image-to-recruitment platform for peripheral nerve stimulation, companion manuscript (in preparation).

[2] E.D. Musselman, J.A. Cariello, W.M. Grill, N.A. Pelot, ASCENT (Automated Simulations to Characterize Electrical Nerve Thresholds): A pipeline for sample-specific computational modeling of electrical stimulation of peripheral nerves, PLOS Computational Biology 17 (2021) e1009285.

[3] T. Couppey, L. Regnacq, R. Giraud, O. Romain, Y. Bornat, F. Kolbl, NRV: an open framework for in silico evaluation of peripheral nerve electrical stimulation strategies, PLOS Computational Biology 20 (2024) e1011826.

[4] I.A. Baratta, et al., DOLFINx: the next generation FEniCS problem solving environment, 2023.

[5] M.L. Hines, N.T. Carnevale, The NEURON simulation environment, Neural Computation 9 (1997) 1179–1209.

[6] M.A. Hussain, W.M. Grill, N.A. Pelot, Highly efficient modeling and optimization of neural fiber responses to electrical stimulation, Nature Communications 15 (2024) 7597.

[7] D.P. Marshall, et al., PyFibers: an open-source NEURON-Python package to simulate responses of model nerve fibers to electrical stimulation, PLOS Computational Biology 21 (12) (2025) e1013764.

[8] C.C. McIntyre, A.G. Richardson, W.M. Grill, Modeling the excitability of mammalian nerve fibers: influence of afterpotentials on the recovery cycle, Journal of Neurophysiology 87 (2002) 995–1006.

[9] D.R. McNeal, Analysis of a model for excitation of myelinated nerve, IEEE Transactions on Biomedical Engineering 23 (1976) 329–337.

[10] N.A. Pelot, B.J. Thio, W.M. Grill, On the parameters used in finite element modeling of compound peripheral nerves, Journal of Neural Engineering 16 (2019) 016007.

[11] C. Geuzaine, J.-F. Remacle, Gmsh: A 3-D finite element mesh generator with built-in pre- and post-processing facilities, International Journal for Numerical Methods in Engineering 79 (2009) 1309–1331.

[12] N. Ravi, et al., SAM 2: Segment Anything in Images and Videos, 2024.

[13] J. Ma, Y. He, F. Li, L. Han, C. You, B. Wang, Segment anything in medical images, Nature Communications 15 (2024) 654.

[14] M. Haberbusch, golgi: an open platform for image-to-recruitment modeling of peripheral nerve stimulation (v1.0.0) [software], Zenodo, 2026. 10.5281/zenodo.21281595.

[15] M. Haberbusch, Reproduction bundles for the golgi peripheral nerve stimulation modeling platform [dataset], Zenodo, 2026. 10.5281/zenodo.21000095.

[16] M. Haberbusch, D. Lung, Y. Jia, R. Blumer, L.F. Reissig, L.M. Zopf, P. Heimel, Rabbit cervical vagus nerve micro-computed tomography dataset from nodose ganglion to superior cardiac branch with tissue segmentations and 3D reconstruction [dataset], Zenodo, 2026. 10.5281/zenodo.21006413.

[17] M. Haberbusch, L.M. Zopf, P. Heimel, R. Blumer, L.F. Reissig, Human cervical vagus nerve µCT dataset from nodose ganglion to superior cardiac branch with tissue segmentations and 3D reconstruction [dataset], Zenodo, 2026. 10.5281/zenodo.21007683.

